# Spatial and temporal assessment of the risk associated to bacteria in recreational waters of a large South American Reservoir

**DOI:** 10.1101/2021.03.15.435485

**Authors:** Daniela Gangi, Diego Frau, Andrea A. Drozd, Facundo Bordet, Soledad Andrade, Mariel Bazzalo, Paula de Tezanos Pinto

## Abstract

The characterization of risk due to recreational exposure to water needs to assess the concentration of pathogens in the water and the degree of contact with those pathogens. In this study we assessed the risk associated to *E. coli* and cyanobacteria in a large South American Reservoir heavily used for recreation, by i) gathering field environmental data from two water agencies (six sites, summers 2011-2015), ii) generating satellite data at landscape scale (750 km^2^, summers 2011-2017) and running a health survey related to water exposure (summer 2017). Field data showed that cyanobacteria abundances recurrently surpassed the moderate and high-risk categories across sites and year analyzed, and a significant positive link between cyanobacteria abundance and microcystin concentration. Nevertheless, microcystin concentrations were in 90% of cases mostly within the low to moderate risk categories. Mean *E. coli* concentrations during 2011-2015 were within the high-risk category in 30% of the sites, but in 2017, sites identified as low risk had high-risk. The latter underscores the high risk posed by *E. coli* in the reservoir. Cyanobacteria (cell abundance and microcystin) and coliform bacteria abundances were unrelated, suggesting different responses to environmental or anthropogenic triggers. Satellite data evidenced that the highest risk related to cyanobacteria abundance occurred in the dendritic areas of the Argentinean side of the reservoir, areas which currently remain unmonitored by water agencies. Satellite monitoring bridged the limited spatial and temporal coverage of field samplings for cyanobacteria abundance (yet not for toxicity nor *E. coli* abundances) and rendered a risk map at landscape scale, which can be used by water agencies to effectively monitor and manage cyanobacteria blooms, and to-coupled with exposure variables-assess health risks related to cyanobacteria. The health survey identified few numbers of suspected patients with symptoms and who bathed in the Salto Grande reservoir. At the time of exposure, sites in the environment evidenced high bacteria concentration (mostly *E. coli* and to a lesser extent cyanobacteria) denoting situations where aspects of the biophysical environment affect human health. More studies and integration among environmental and health disciplines are needed to assess the impacts of these water born bacteria in human health. Finally, we further assessed how well cyanobacteria quantitative proxies monitored in the field explained the outcome of a qualitative risk communication system-the cyano-traffic-light-which is ongoing since 2011. We obtained a significant predictive model only for cyanobacteria abundance, yet with low predictive value. This probably occurred because the variables used to build each cyano-traffic-light category (cyanobacteria abundance, toxicity and chlorophyll-a, scums) were monitored with different frequencies, and because at least two of these variables needed to surpass the threshold of each category to be allocated into a risk category. Based upon our results we propose several modifications to the current cyano-traffic-light, that believe would better reflect what happens in the field and protect human health: i) include *E. coli* concentration and satellite estimated cyanobacteria abundance (mostly in areas not covered by field monitoring), ii) relax the thresholds for cyanobacteria abundance and toxicity, and iv) base each risk category upon the surpassing of one of either *E. coli,* cyanobacteria abundance, microcystin.

## Introduction

Recreational coastal water and freshwater environments are areas where leisure activities are undertaken by end users. The contact with water in recreational areas can be direct through sports, like swimming, surfing, and diving, or indirect, such as fishing, sailing, walking, and picnicking. Health risks associated with water recreation may be caused by infectious microorganisms (bacteria, virus, and protozoa), geogenic substances (arsenic, etc.), industrial and agricultural chemicals (pesticides etc.) and toxins produced by cyanobacteria (Chorus and Welker, 2021).

Among bacteria hazards in water, Cyanobacteria, which are not in themselves infectious, have the capacity to produce hepatotoxins and neurotoxins that can affect human health (Stewart et al. 2006; Wood 2016). Cyanotoxins produced by cyanobacteria constitute a global water-quality concern (Chorus and Bartram, 1999; Paerl and Scott, 2010). Though several studies show that cyanobacteria abundance is positively linked to their toxin concentration (Pearl and Otten 2013; Ibelings et al. 2014; Bordet et al. 2017), high cyanobacteria abundances can occur at low or without toxin production (Paerl and Otten 2013). Potential exposure routes to cyanobacteria and its cyanotoxins may include ingestion, contact, or inhalation during recreational activities (Chorus and Bartram, 1999), and may cause health effects such as vomits, diarrhea and headaches (Chorus et al.2000; Halac et al. 2019), skin reactions (Pilotto et al., 2004; Zanchett and Oliveira-Filho, 2013) and respiratory symptoms (Stewart et al. 2006a). Also, toxins may cause indirect health effects due to the consumption of fish containing high toxin concentrations (Freitas de Magalhães et al. 2001; Giliane and Oliveira-Filho, 2013). Finally, the study of Stewart et al. (2006a) also found higher respiratory symptoms at recreational sites with high cyanobacteria abundance yet without toxin concentration, compared to sites with low cyanobacteria abundance.

Coliform bacteria in recreational water indicates that fecal pollution may have occurred, and pathogens might be present (Tallon et al. 2005; Boehm et al, 2009). Indeed, sewage water contamination produces between 6 to 60 billion cases of gastrointestinal illness annually (Caslake et al., 2004) and respiratory, eye, ear, nose, throat, and skin infections (Russo et al. 2020). Many of these symptoms are like the ones caused by recreational exposure to cyanobacteria.

Evidence in the literature, suggest that pathogens different from cyanobacteria may coincide during cyanobacteria blooms. For example, several studies showed that during cyanobacteria blooms, heterotrophic bacteria concentration was higher than in situations without cyanobacteria blooms (Bouvy et al. 2001; Eiler and Bertilsson 2007; Abdulaziz et al. 2016; Wang et al. 2016). Also, Berg et al. (2009) found positive effects of cyanobacteria on bacteria probably due to the presence of organic substances in cyanobacteria biofilm. On the contrary, compounds present in some cyanobacteria species (e.g., *Microcystis aeruginosa*) have been reported to affect the survival and growth of some bacteria species (Østensvik et al. 1998; Skulberg 2000; Bomo et al. 2011). We could not find a study assessing the link between cyanobacteria and sewage bacteria, yet in many water bodies both hazards coincide (Halac et al. 2019).

Recreational water quality guidelines protect the public from health risks associated with water recreation (Russo et al. 2020). For Cyanobacteria, the World Health Organization (WHO) established three risk categories for recreative use of water, based upon: the abundance of cyanobacteria, the concentration of chlorophyll-a and the presence of scums at the water surface (Chorus et al., 2000; Chorus, 2012, Chorus and Welker 2021). A given risk category is established upon the surpassing of at least one of the mentioned variables. Likewise, for coliform bacteria, standards for recreational water quality had been developed by several agencies, as for example the Environmental Protection Agency (EPA) and the European Directive for Recreational Water (Water Framework Directive, WFD). For both cyanobacteria and coliform bacteria, depending on the risk category, risk can be low, moderate, and high and this has implications (allowed, banned) for safe use of water for recreation.

A necessary step towards preventing, anticipating, and mitigating the risk posed by cyanobacteria and coliform bacteria on human health is to characterize their concentration in the environment, both in space and time. Therefore, permanent monitoring of waterbodies at large spatial and temporal scales is needed. For cyanobacteria monitoring, traditional methods include collection of water samples followed by microscopic identification and quantification. Field monitoring approaches are time and labor intensive, and, despite the effort and costs involved, they often fail to represent the spatial and temporal heterogeneity of cyanobacterial blooms, which are very dynamic, particularly in large ecosystems. More recently, cyanobacteria blooms are monitored using remote sensing techniques, by estimating several cyanobacteria proxies such as chlorophyll-a (Tebbs et al. 2013; for a review, see Dörnhöfer and Oppelt 2016), phycocyanin (Simis et al. 2005; Randolph et al. 2008; Li et al. 2015) or cell numbers (Lunetta et al. 2015; Drozd et al. 2019). Indeed, remote sensing is recognized as a tool for providing complete and synoptic geographical coverage of water quality in freshwater systems (Hadjimitsis et al. 2010 and references therein). Regarding cyanobacteria blooms, the use of satellites allows assessing bloom extent (Kutser 2004; Alikas et al. 2010), frequency (Kahru et al. 2007) and temporal trends (Stumpf et al. 2012, Urquhart et al. 2017). This information can complement the information obtained in field monitoring programs (Klemas 2012) and help to assess risk. Indeed, the near real time availability of water quality data available from current satellites makes it possible to integrate satellite data into early warning systems to protect human health and ecosystems (Welker et al. 2021). Nevertheless, cyanobacteria toxins (cyanotoxins) and coliform bacteria are optically inactive and hence cannot be monitored using satellite images. Paradoxically, despite of the great utility of the use of satellites in the assessment of water quality, still few studies use these tools compared to studies performed exclusively in the field, as shown by Dörnhöfer and Oppelt (2016).

Illness risk is associated with both the concentration of pathogens in the water and the degree of contact with those pathogens (Russo et al. 2020). Evidence in the literature of illness due to recreational exposure to water, however, is scarce; for example, the literature review performed by Russo et al (2020) only found 112 studies which reported (at least) a health symptom associated to ambient surface water exposure. For cyanobacteria, despite a wealth of studies regarding the ecology and toxicity of cyanobacteria, the number of epidemiological studies of health effects due to cyanobacteria exposure is scarce, as evidenced by the reviews of Stewart et al. (2006b) and Wood (2016). Suspected health cases of cyanobacteria intoxication include those where there is evidence of recreational use of water containing cyanobacteria and where the symptoms cannot be linked to another agent (Wood 2016). Identifying illness due to cyanobacteria exposure is challenging, as other microbial agents (*E. coli*, gram negative bacteria, amoeba, and viruses) present in the water may cause symptoms like those of cyanobacteria (Stewart et al. 2006b).

In South America, the Salto Grande Reservoir (Uruguay River), suffers recurrent and massive cyanobacteria blooms (Bordet 2009; Chalar 2009; O’Farrell et al. 2012; Bordet et al. 2017; Gangi et al. 2020), has high concentration of coliform bacteria (CARU 2016, Rougier et al 2018), and is heavily used for recreation during the warm season. This area has been monitored by two water agencies - *Comisión Administradora del Río Uruguay* (CARU) and the *Comisión Técnica Mixta* (CTM)-for about two decades. Since 2011 a local risk communication system for recreational use of the waters was implemented by CARU. This system is called the cyano-traffic-light as it informs the risk posed by cyanobacteria following the colors of a traffic light (red bans bathing, yellow suggests avoiding bathing and green allows bathing). The quantitative proxies and thresholds used to build the cyano-traffic-light include cyanobacteria abundance, chlorophyll-a concentration, scum formation and cyanotoxin concentration. Risk thresholds of chlorophyll-a and presence of scum are based upon the guidelines established by WHO, while risk thresholds for cyanobacteria are based upon the guidelines established by Australia (which are more conservative than those of the WHO). In the cyano-traffic-light the establishment of a given risk category is determined by the surpassing of at least two of the four proxies described above. Despite the cyano-traffic-light has been used for about 10 years with weekly or fortnightly temporal resolution, up to date it is unclear how well the quantitative variables used to build the cyano-traffic-light explain the outcome of these reports. Moreover, up to date reports are presented in isolated formats and have not yet been synthesized for assessing annual or interannual trends, albeit for short periods (Rougier et al. 2018). Finally, in the Salto Grande Reservoir, despite a wealth of environmental information available for cyanobacteria, and to a lesser extent for coliform bacteria (Bordet 2009; Chalar 2009; O’Farrell et al. 2012; CARU 2016; Bordet et al. 2017; Rougier et al. 2018; Gangi et al. 2020), there is a remarkable absence of documentation of human illness that could be related to recreational exposure, except for one case of a young boy who was exposed to a dense bloom of cyanobacteria while using his jet ski and who developed serious health effects (Giannuzzi et al. 2011).

In this study, we assessed the spatial and temporal distribution of the risk due to cyanobacteria and *E. coli*, and its potential link to human illness due to recreative exposure, in the Salto Grande Reservoir (Uruguay River). To achieve this, we integrated different approaches, scales, and sources of information: 1). First, at field scale, along a five-year period (2011-2015), we analyzed cyanobacteria and *E. coli* data from water at six recreational areas, using available quantitative (cyanobacteria abundance, toxicity, and coliform bacteria abundance; chlorophyll-a data were unfortunately unavailable) and qualitative (cyano-traffic light reports) data from the two water agencies that monitor the reservoir. With this information we: i) characterized the spatial and temporal distribution of these hazardous bacteria, ii) explored the relation among bacteria proxies (cyanobacteria and coliform bacteria, cyanobacteria and cyanotoxins, and coliform bacteria and cyanotoxins) and iii) assessed how well quantitative cyanobacteria proxies explained the outcome of the risk communication system (cyano-traffic-light). 2) Second, at landscape scale, using satellite images, we expanded the time frame available (2011-2017) and spatial scale (750 km^2^) for the variables of cyanobacteria abundance and chlorophyll-a. 3) Third, in summer 2017, a survey was run by physicians and health assistants in the largest hospital of the area, for identifying cases of symptoms that could be related to cyanobacteria and *E. coli* and who had used the reservoir for recreation. Finally, we assessed the risk posed by these bacteria at the site and time of recreational exposure.

## Material and Methods

### Study Area

The Salto Grande Reservoir (29.43-31.12° S, 57.06-57.55° W) is a large reservoir (750 km^2^) located in the lower stretch of the Uruguay River, a natural limit between Argentina and Uruguay, in South America. This reservoir has a river like main channel and five lateral arms with a dendritic morphology where 12 public recreational areas are located (Fig. 1). These public recreational areas have amenities and lifeguard, yet many other unofficial recreational areas can be accessed by the public and landowners along its coastline. Energy production is the main ecosystem services in the area; cultural and recreational services (bathing, water sports, and fishing) are also important, mostly during summertime. Populations in the largest cities located close to the riverside (e.g., Concordia city 149,450 inhabitants and Federación city 16,658 inhabitants. Source: INDEC 2010) significantly increase during summer due to the tourism (7 and 400%, for Concordia and Federación, respectively). This reservoir is eutrophic (Bordet et al. 2017) and has been suffering massive and recurrent cyanobacteria blooms during the last 20 years (Bordet 2009, O’Farrell et al. 2012, CARU 2016, Bordet et al. 2017, Izaguirre et al. 2018; Rougier et al. 2018, Gangi et al., 2020). Cyanobacteria blooms occur mostly in summer-coinciding with the peak tourism season- and in autumn. Cyanobacteria blooms are mostly dominated by *Microcystis aeruginosa*, and to a lesser extent by several species of the genera *Dolichospermum* (Bordet et al. 2017); these cyanobacteria can produce the hepatotoxin microcystin. Bordet et al. (2017) and Martinez de la Escalera et al. (2017) recorded microcystins in the reservoir, at a wide range of concentrations (0-621 µg L^-1^). Moreover, there is evidence of coliform bacteria at thresholds above the guidelines of safe use of recreational water in different locations of the reservoir (CARU 2016; Rougier et al. 2018).

**Figure. 1.**
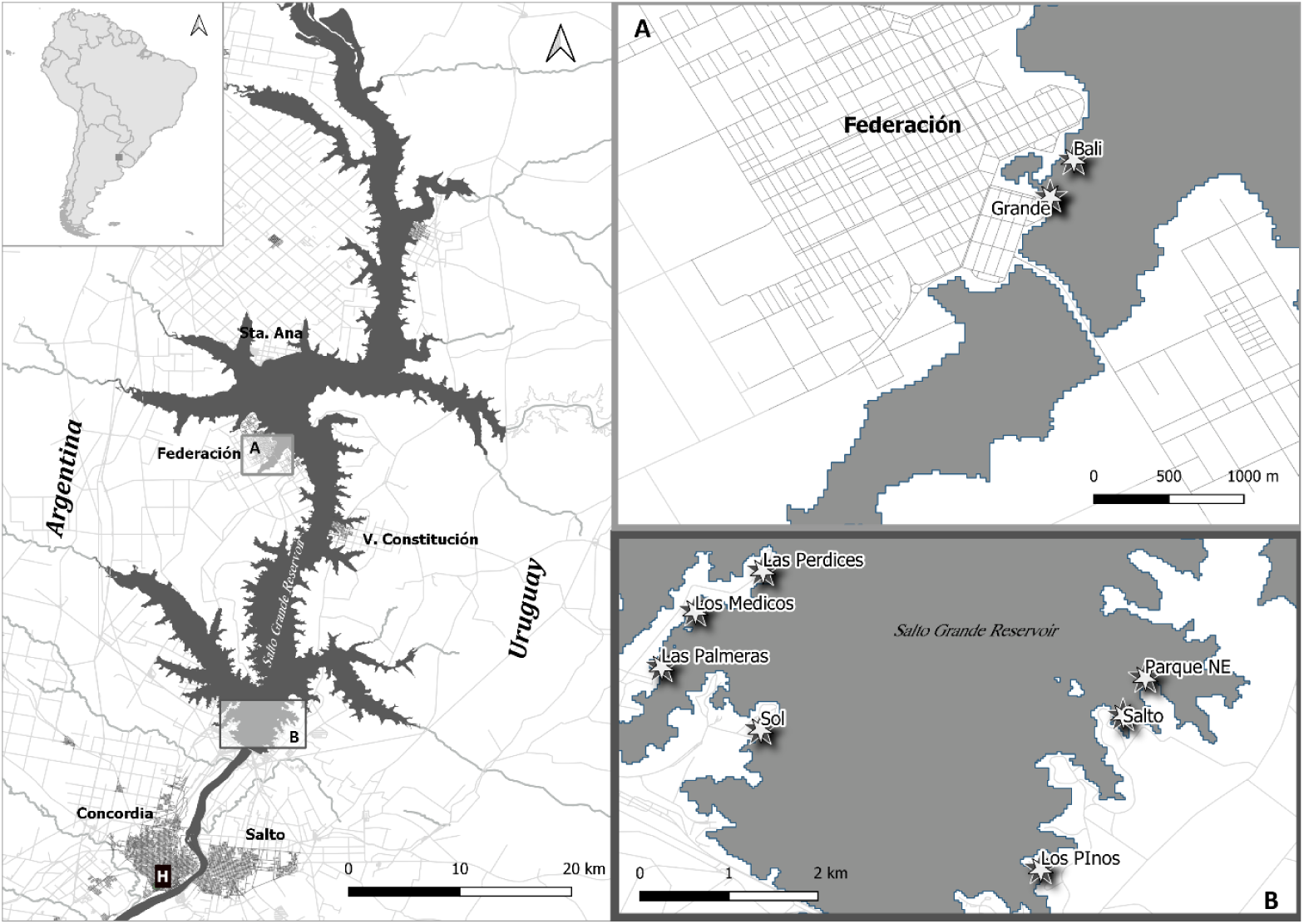
Location of the Salto Grande Reservoir (Uruguay River). Subsets A and B show the six recreational areas where water was sampled (A=sites located in the northern part, Federación city, Subset B= sites located in the southern part). H denotes the location of the biggest hospital of the area, where the survey was performed, which is located 15km away from the recreational areas in subset B.

### Data collection and analyses

The spatial and temporal distribution of the risk posed by cyanobacteria and *E. coli*, and its potential link to human illness due to recreative exposure, in the Salto Grande Reservoir (Uruguay River) was assessed using 3 different sources of information:

### Assessment of the spatial and temporal distribution of cyanobacteria and coliform bacteria: water monitoring at recreational areas

We worked with six recreational areas distributed along the Argentinean side of the Salto Grande Reservoir: two located in the city of Federación and four located 15km up north from the city of Concordia (Fig 1). For the six recreational sites, we gathered all available information from two water agencies, the Comisión Técnica Mixta Salto Grande (CTM) and Comisión Administradora del Río Uruguay (CARU), within the period December to March (where there is coincidence between cyanobacteria blooms and the tourism season) of the years 2011-2017. Data for 2016 and 2017 were incomplete and hence could not be used for running the analyses, and the year 2014 was removed from the analysis because of absence of cyanobacteria blooms. The quantitative data used to run the analyses included: cyanobacteria abundance (cell ml^-1^) (n= 213), semi-quantitative concentration of microcystins (µg L^-1^) (n= 76), and total coliform concentration (CFU 100 mL^-1^) (n= 209). Note that the monitoring of toxicity was less frequent than the monitoring of cyanobacteria abundance. Unfortunately, chlorophyll-a concentrations were only available for the December to March of the year 2015 (n= 10) (Izaguirre et al. 2018) and hence were not used to compare spatio-temporal scales but was used in the PCA and in the predictive model (see below).

Cyanobacteria cyanotoxin (microcystin) and chlorophyll-a samples were collected using plastic flasks and submerging them below 0.3 meters from the surface. Samples for cyanobacteria quantification were immediately fixed with Lugol 1% acidified solution and counted in inverse microscope following Utermöhl (1958). When an extremely high abundance was detected, the counting was performed using a Neubauer haemocytometer on an optical microscope after hot sodium hydroxide digestion (Reynolds and Jaworski 1978). For microcystin estimation, 20 mL of unfiltered sample was exposed to 3 cycles of freezing and thawing and analyzed with the Abraxis®13 semi-quantitative method that detects concentrations within the following categories: 1; 2,5; 5; 10 and >10 µg L^-1^). The samples for chlorophyll-a were stored in dark and cold conditions until analyzed in the laboratory, and processed following standard methods (APHA, 2012). Fecal coliforms were obtained using sterile plastic flasks and collecting water samples at 0.3 m below the surface. The samples were immediately stored in cold and dark conditions, and estimated following the 9222D standard method (APHA, 2012). Next, *Escherichia coli* was estimated from fecal coliforms using a conversion factor of 0.77 (Rasmussen and Ziegler, 2003) and used for assessing health risks in recreational waters.

The risk posed by cyanobacteria in recreational waters was assessed following tow guidelines: World Health Organization (WHO) and cyano-traffic-light developed by the Comisión Administradora del Río Uruguay (CARU) (Table 1). The risk posed by *E. coli* in recreational waters was assessed using the directives established by the Environmental Protection Agency (EPA, 500 CFU 100mL^-1^) and the European Directive for recreational use of water (EU, 900 CFU 100mL^-1^).

**Table 1.** Criteria used by the Comisión Administradora del Río Uruguay (CARU) for establishing the risk categories of the cyano-traffic-light for recreative use of water, the colors assigned for each risk category and the recommendation for recreational use of waters.

| Proxy | Low Risk | Moderate Risk | High Risk |
| --- | --- | --- | --- |
| Chlorophyll-a ($\mu\text{g L}^{-1}$) | <10 | 10-50 | >50 |
| Cyanobacteria (cell mL<sup>-1</sup>) | <5,000 | 5,000-50,000 | >50,000 |
| Microcystin ($\mu\text{g L}^{-1}$) | < 2 | 2-10 | > 10 |
| Scums at surface | Absence | Absence | Presence |
| Cyano-traffic-light color | Green | Yellow | Red |
| Bathing recommendation | Allowed | Avoid using | Prohibition of use |

Note that for chlorophyll-a concentration and scum presence, the criteria of the World Health Organization (WHO) and CARU coincide, whereas for cyanobacteria abundance thresholds in WHO are less conservative (low risk: <20,000 cell/ml, moderate risk <100,000 cell/ml, high risk: >100,000 cell/ml) than CARU.

For testing differences among the quantitative variables (cyanobacteria abundance, cyanobacteria toxins and *E. coli*), recreational areas (six sites) and years with sufficient information (2011, 2012, 2013, 2015), we run Generalized Linear Models (GLM) with Gaussian adjustment. Bonferroni post-tests were used to compare the pair-samples for beaches and years. All data used were transformed to the log10 (x+1) for the analyses.

We used contingency tables (Supplementary Material 1) to explore the possible association between *E. coli* and cyanobacteria abundance, between *E. coli* abundance and microcystin concentration, and between cyanobacteria abundance and microcystin concentration. For this, we constructed categories based on international standards for recreational water use (Supplementary Material 1). The Fisher’s exact test was used to determine statistically significant correlations among categories of the variables, and the Cramer’s V test was used to determine the correlation level, from lack of association (0) to complete association (1) between two variables.

Moreover, we assessed the spatial and temporal distribution of the cyano-traffic-light reports. These reports provide qualitative information regarding the risk posed by recreational exposure to cyanobacteria using the colors of a traffic light: high risk (red color, bathing is banned), moderate risk (yellow color, bathing should be avoided) and no risk (green color, bating is allowed) (Table 1). The risk thresholds informed in each report are computed based on four criteria: chlorophyll-a concentration, cyanobacterial cell abundance, microcystin concentration and presence of scums at the water surface (Table 1). At least two criteria must be accomplished to establish a given risk category, yet this information is not explicitly provided in the reports, and sometimes expert assessment is used to establish the informed risk category. We downloaded all available reports from the official website of CARU (http://www.caru.org.uy/web/) within the warm period (December to March) of the years (2011, 2012, 2013, 2015), following the same spatial and temporal scale than the one available for quantitative variables. For each year and recreational site analyzed we assessed the number of times each of the 3 categories (low, moderate, and high risk) was recorded. Next, to test how well quantitative variables related to cyanobacteria (cell abundance, toxin concentration, chlorophyll-a) explained the outcome of the cyano-traffic-light, we ran multiple regression models with Gaussian adjustment. We used the following quantitative variables as predictors: cyanobacteria abundance (n= 64), microcystin concentrations (n= 22) and chlorophyll-a concentration (n= 10). The observed and statistically significant models were compared by using a dispersion plot.

### Assessment of the spatial and temporal distribution of the risk posed by cyanobacteria at landscape scale: remote sensor monitoring

To obtain mean values for the variables of cyanobacteria abundance and chlorophyll-a for the warm period (December to March) of the years 2011-2017, we processed Level 2 Landsat 7 ETM+ and Landsat 8 OLI images, at surface reflectance in Google Earth Engine platform (Gorelick et al. 2017). We processed a total of 76 images; note that 2 images needed to be processed per sampling date to cover the entire reservoir. The images were first filtered with a total cloud cover less than 15%, then we selected the water body of Salto Grande by a threshold of 0.025 on SWIR band that worked also as a cloud mask. To prevent confusions between phytoplankton and suspended matter, we also masked pixels where red band reflectance was higher or equal to the green band. In those pixels where the green band was higher than the red band, we computed cyanobacteria abundance and chlorophyll-a concentrations using algorithms built using spectral firms obtained in the study area by Drozd et al. (2019) (Table 2).

**Table 2.** Algorithms used for estimating chlorophyll-a concentration and cyanobacteria abundance using Landsat satellites, developed by Drozd et al. (2019) for the Salto Grande Reservoir. λg=green band, λr=red band and λnir=near infra-red band.

| Satellite | Chlorophyll-a concentration | Cyanobacteria abundance |
| --- | --- | --- |
| Landsat 7 ETM+ | $e^{1.96+8.51*x}$ | $e^{4.35+28.8*x-19.4*x^2}$ |
| Landsat 8 OLI | $e^{2.2+9.55*x}$ | $e^{5.35+26.25*x-17.38*x^2}$ |
| Algorithm | $\frac{(\lambda g - \lambda r + \lambda nir)}{(\lambda g + \lambda r + \lambda nir)}$ | |

The cyanobacteria abundance algorithm can detect average densities of 200 cells mL^-1^ (Drozd et al. 2019) and hence is able to map CARU’s alert levels. Next, for the mean values obtained during the studied period we classified pixels into the alert categories proposed by WHO and CARU, calculated the areas of each risk category and built maps showing of the distribution of the risk at landscape scale (750 km^2^).

To explore the relation among variables assessed using field and satellite metrics, we run a principal component analysis (PCA) whenever field and satellites data were obtained at the same day or up to one week apart (n= 25). All data were Log10 transformed. For field data the following variables were used: cyanobacteria abundance, chlorophyll-a, and the cyano-traffic-light (microcystin data were too scares to be used). The qualitative variable cyano-traffic-light was computed as a quantitative variable by considering each report category as follows: 1=low risk, 2=moderate risk, 3= high risk. For satellite data the following variables were used: cyanobacteria abundance and chlorophyll-a.

### Assessment of cases of possible human intoxication due to recreational exposure to hazardous bacteria

In summer (January and February) 2017, a survey was performed to identify suspected clinical cases with health symptoms associated to bacteria exposure (gastrointestinal, eye, ear, dermal illness) and who had recreationally exposed to waters along the Salto Grande Reservoir. The survey was collected in the emergency room of the Masvernat Hospital, the largest hospital in the area, located 15 km south from the reservoir (Fig 1). The implementation of the study involved different preparation phases including: an ethical evaluation of the project, signing a contract for information exchange between the hospital and academia, holding a meeting with physicians and health assistants at the hospital, designing the survey and establishing the working methodology. The survey was based upon the revision of publications (Stewart 2004; Stewart et al. 2006a) and upon a shorter version of the survey previously tested in the Masvernat Hospital. The survey gathered information regarding if recreational activity was undertaken in the Salto Grande Reservoir, location, date, type (swimming, fishing) and duration (half hour, one hour, two hours) of the recreational activity, symptoms (gastrointestinal, eye, ear, dermal illness), time lapse between exposure and onset of symptoms, and demographic data (Supplementary material 2). The study population comprised adults and children who visited the emergency room of the Masvernat Hospitals with symptoms and who had engaged in recreational activities in waters of the Salto Grande Reservoir. The survey was fulfilled by health assistants after a physician determined that the patient presented symptoms. To avoid influencing responses, neither the physician nor the health assistant informed patients about the scope of the study until after the survey was fulfilled. Note that cases with symptoms were considered suspected cases (but not confirmed cases) as serological tests for cyanotoxins or freshwater microbial agents were not available. Cyanobacteria and coliform related illness were considered whenever individuals presenting symptoms had exposed to freshwater contact containing high concentrations of these agents in the site and time of recreation. While other freshwater microbial agents such as gram-negative bacteria, amoeba, and virus may also cause similar symptoms (gastrointestinal, eye, ear or dermal) their influence could not be assessed in this study as they are not monitored in field studies. Finally, for each suspected case we collected environmental data of cyanobacteria abundance, microcystin concentration, coliform bacteria concentration, and cyano-traffic-light category, from the weekly monitoring program run by CTM and CARU, for the field and time of exposure. When the date of field sampling did not coincide with the date of recreational exposure, we used the environmental data from the closest date available.

## Results

### Assessment of the spatial and temporal distribution of cyanobacteria and coliform bacteria: water monitoring at recreational areas

Cyanobacteria mean abundance showed a high spatial and temporal heterogeneity (Fig. 2a), with statistical different abundances among recreational sites (Wald-Chi Square= 23.95, P< 0.001), years (Wald-Chi Square= 154.42, P< 0.001) and by the effect of interaction of both factors (Wald-Chi Square= 183.42, P< 0.001). In 2011, five (Las Perdices, Los Medicos, Grande and Baly) of the six recreational sites analyzed had mean cyanobacteria densities that surpassed the highest risk categories of the Comisión Administradora del Río Uruguay (CARU high risk >50,000 cells mL^-1^), and two of these sites (Las Perdices and Los Medicos) also surpassed the highest risk level established by the World Health Organization (WHO Alert level II >100,000 cells mL^-1^) (Fig 2a). Compared to the other years, 2011 had significantly higher abundance than 2013 and 2015, but significantly lower abundances than 2012 (Table 3). In 2012, all recreational sites analyzed had mean cyanobacteria densities that surpassed the highest risk categories of CARU (>50,000 cells mL^-1^) and WHO (Alert level II >100,000 cells mL^-1^) (Fig 2a). Indeed, the year 2012 had significantly higher abundances than any other year analyzed (Table 3). In 2013 and 2015, cyanobacteria abundances were mostly within the moderate risk category established by CARU (>5,000-<50,000 cells mL^-1^) and within the low-risk category established by the WHO (<20,000 cells mL^-1^) (Fig. 2a). The year 2013 had, nevertheless, significantly higher abundances than 2015 (Table 3). Note that the year 2014 was excluded from the analysis as it lacked cyanobacteria blooms. Differences among recreational sites were significant between Las Palmeras, which had significantly higher cyanobacteria abundance than Las Perdices, Sol and Grande (Table 3). Microcystins were detected in all recreational sites and years analyzed; mean microcystin concentrations were mostly low (≤2 µg L^-1^, 65%), on few occasions moderate (>2<10 µg L^-1^, 25%) and in rare occasions high (>10 µg L^-1^, 10%) (Fig. 2a). The relation between cyanobacteria abundance and microcystin concentration was significant and positive (Fisher’s exact P= 0.01; Cramer’s V= 0.65).

**Figure 2.**
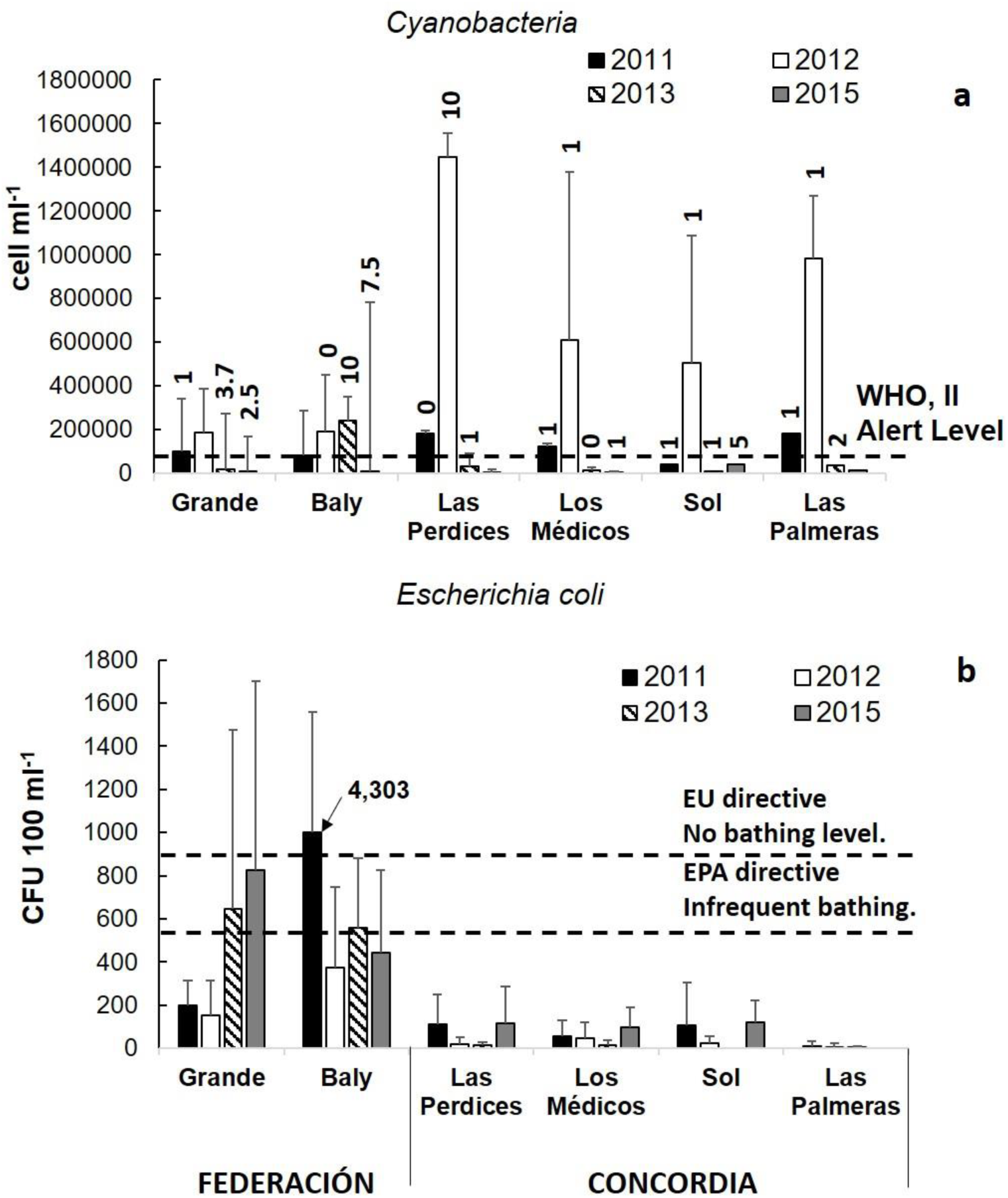
Mean cyanobacteria abundance (cell m L^-1^), mean microcystin concentration (µg L^-1^) (a), and mean *E. coli* abundance (CFU100 m L^-1^) (b) at the six recreational areas and warm periods of the 4 years analyzed. The Alert level II of WHO, >100,000 cells mL^-1^, is depicted in a). Note that the CARU high risk threshold, >50,000 cells mL^-1^, is not shown in a) because of the scale of the graph.

**Table 3.**
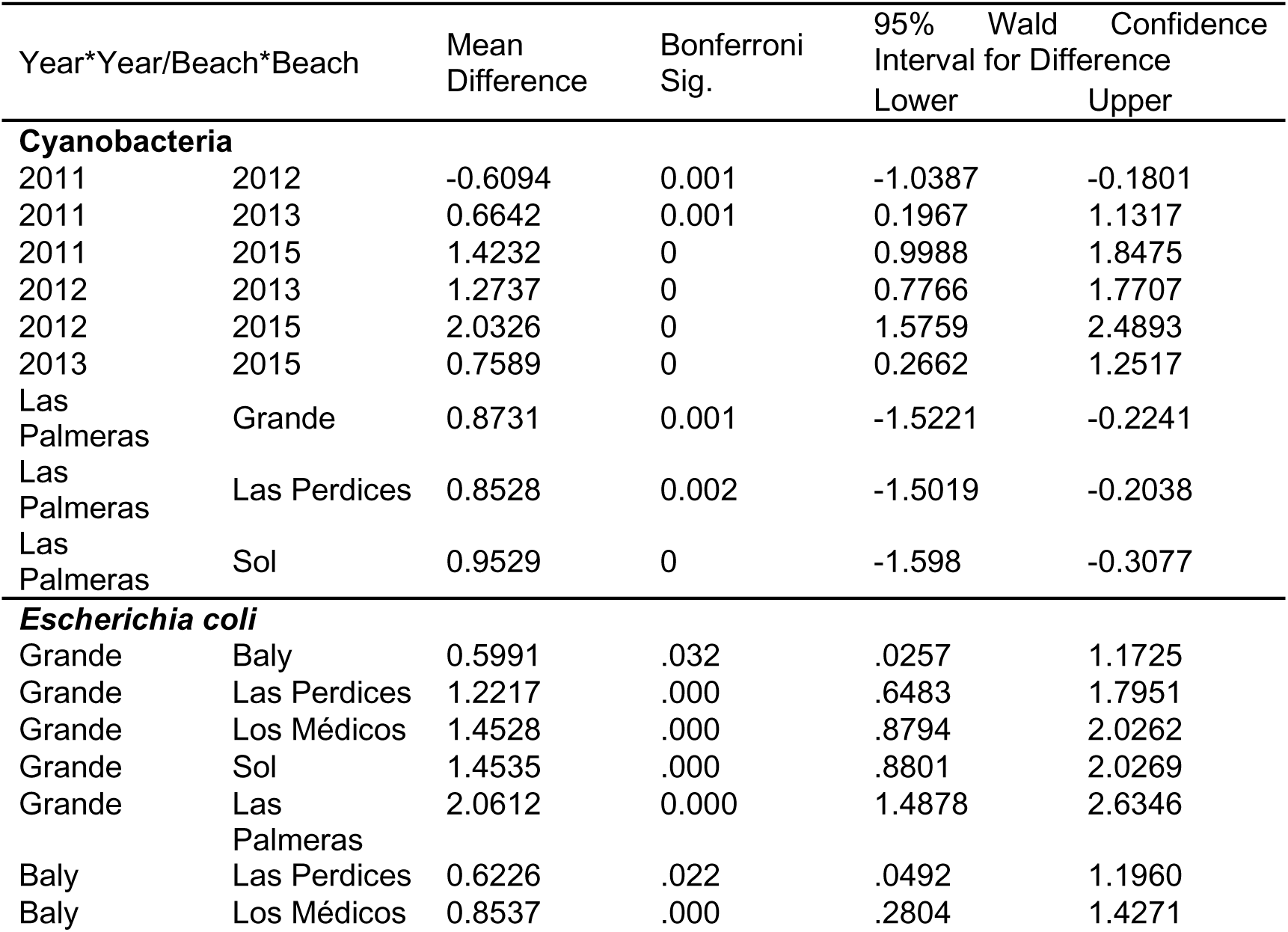
Bonferroni post-hoc comparisons for the variables of cyanobacteria and *E. coli* abundance for the years and sites analyzed in the field and which showed statistical differences (P < 0.05).

Mean *E. coli* values in the recreational sites located in Federación department recurrently surpassed the thresholds for recreational use of water established by the Environmental Protection Agency (EPA) (2002), and in occasions the thresholds of the European Bathing Water Quality Directive (EU) (2006/7/EC) (Fig. 2b). Noteworthy, *E. coli* concentration in the recreational site Baly was extremely high in 2011 (4300 CFU 100ml^-1^) (Fig. 2b). Recreational sites located in Concordia department always had mean *E. coli* concentrations below EU and EPA risk thresholds (Fig. 2b). The GLM analysis showed significant differences in coliform bacteria concentration among recreational sites (Wald-Chi Square= 138.52, P< 0.001, both sites in Federación were significantly higher than sites in Concordia department) and for the interaction of space and time (Wald-Chi Square= 84.38 P< 0.001), yet similar results among years (Wald-Chi Square= 6.59 P= 0.086). Finally, we found absence of relation between *E. coli* abundance and cyanobacteria abundance (Fisher’s exact P= 0.64; Cramer’s V= 0.10), and between *E. coli* abundance and microcystin concentration (Fisher’s exact P= 0.25; Cramer’s V= 0.21).

The cyano-traffic-light reports in the year 2012 were mostly within the high-risk category (54%), whereas in the years 2011, 2013 and 2015 most reports indicated low risk (46, 67 and 50%, respectively) (Fig. 3, Supplementary material 3a). Note that the year 2014 had absence of blooms and hence was not analyzed here. Among recreational sites, the highest percentage of reports within the high-risk category occurred in Perdices, Los Médicos and Palmeras (mean percentage range: 28-34%), located in Concordia department (Supplementary material 3b).

**Figure 3.**
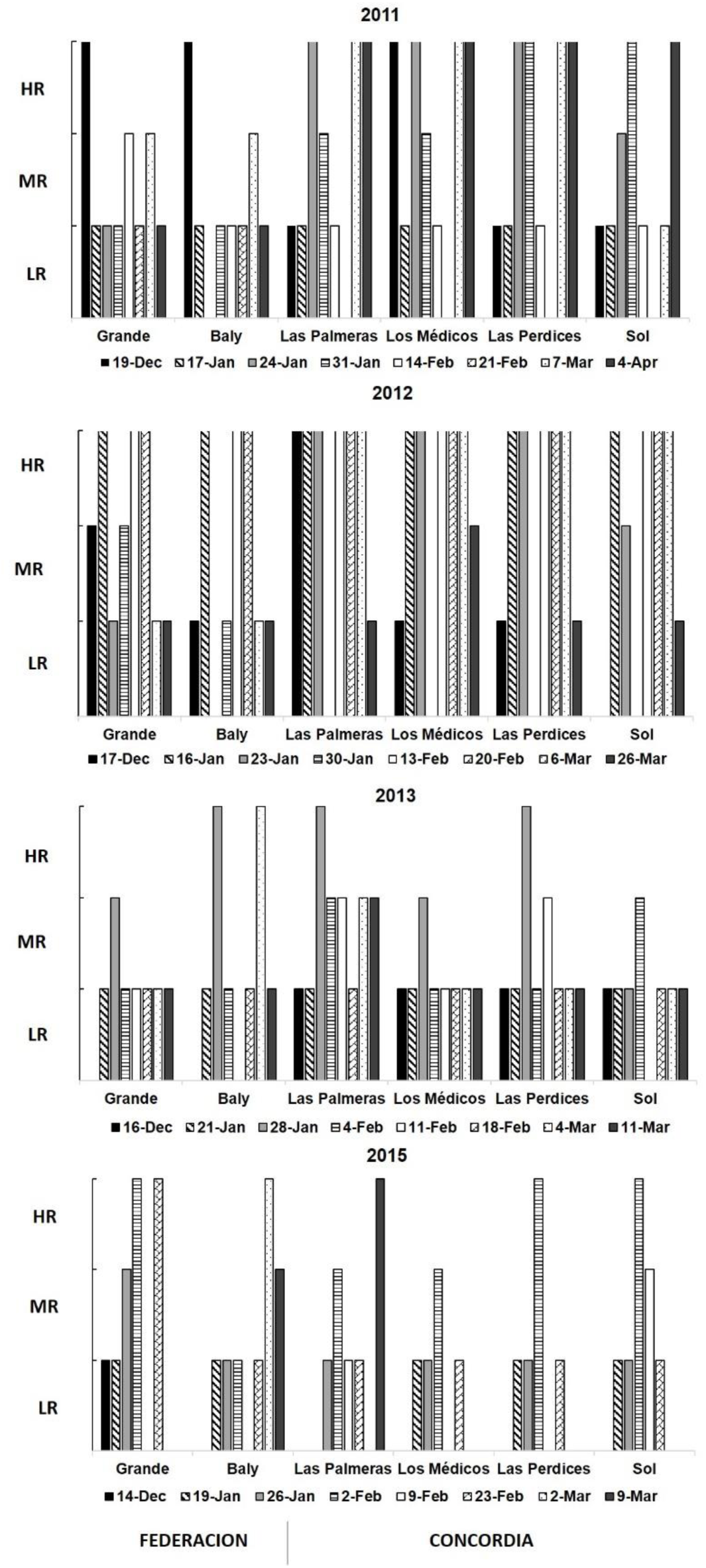
Cyano-traffic-light categories for each (warm period of each) year and each recreational area analyzed (n=159). Low risk (LW), moderate risk (MR), and high risk (HR).

The multiple regression model using cyanobacteria abundance as a predictor of the categories of the cyano-traffic-light was significant (predicted cyano-traffic light= 1.59173 + (cyanobacteria abundance) *0.00000132, F= 43.4, P< 0.001), and explained 40.2% of the total variability. Noteworthy, other models ran using other cyanobacteria proxies (chlorophyll-a, microcystins) were non-significant. Based on the significant model (total cyanobacteria as predictor), the predicted cyano-traffic-light and the observed cyano-traffic-light showed a low level of coincidence (21%, Fig. 4), with the predicted model mostly overestimating (47%) or underestimating (32%) the observed cyano-traffic-light.

**Figure 4.**
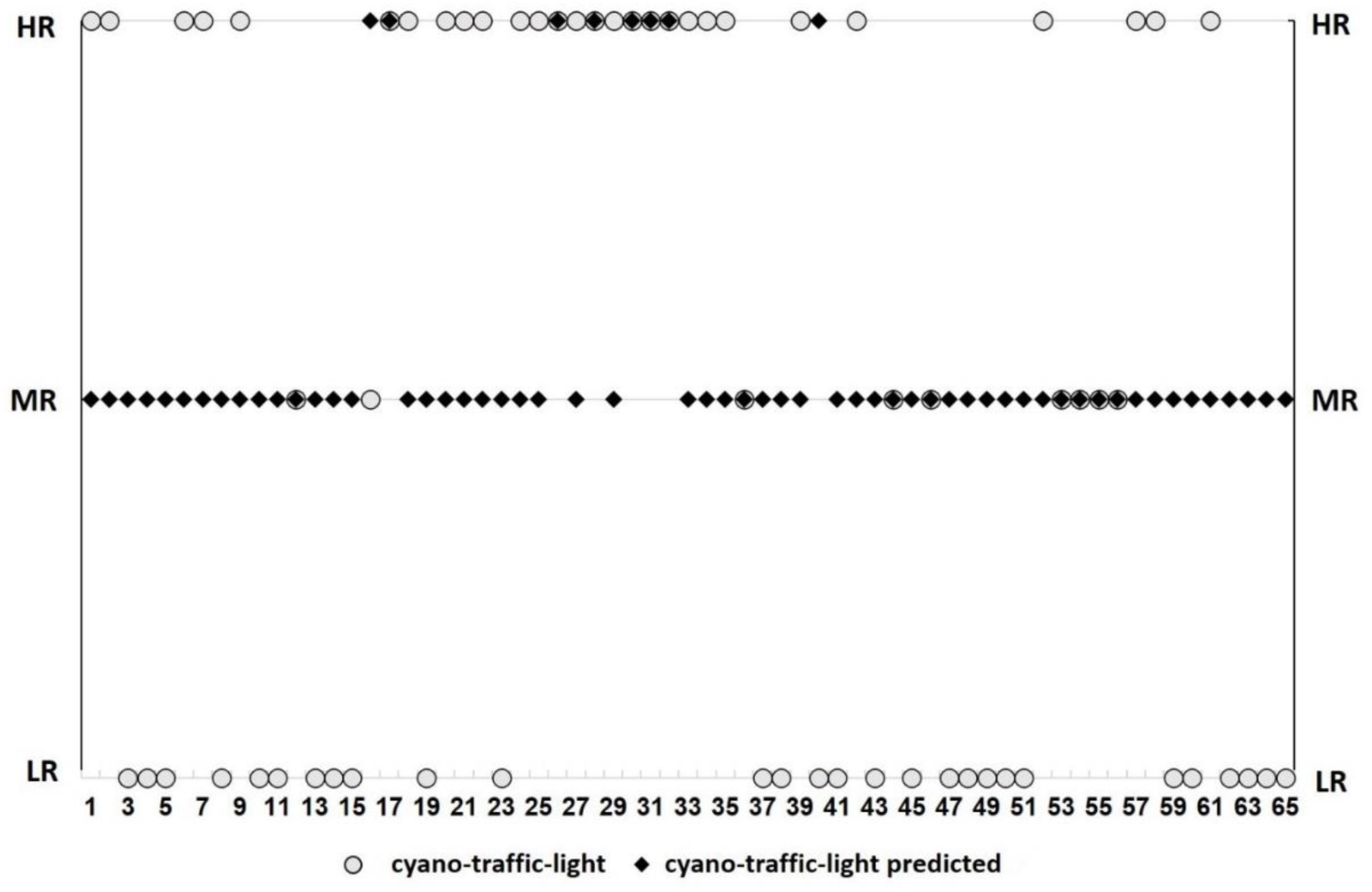
Dispersion plot comparing the results of the predicted cyano-traffic-light using the significant regression model (that used cyanobacteria abundance) and the observed cyano-traffic-light reports. LR= low risk, MR= moderate risk, HR=high risk.

### Assessment of the spatial and temporal distribution of the risk posed by cyanobacteria at landscape scale: remote sensor monitoring

Figure 5 shows the distribution of chlorophyll-a concentration and cyanobacteria abundance at landscape scale (750 km^2^) for the period 2001-2017, classified following the risk thresholds proposed by CARU. Both variables showed a similar distribution: the river-like area showed low risk, whereas arms and the coastline had either low, moderate, or high risk (Fig. 5). The inner part of the arms in the Argentinean side where the sites with higher risk, for both proxies. High risk areas covered from 5909 to 8510 hectares, for cyanobacteria abundance and chlorophyll-a concentration, respectively (13-19% of the reservoir’s surface, respectively). Moderate risk areas covered 11348 to 10675 ha, for cyanobacteria abundance and chlorophyll-a concentration, respectively (25-24% of the reservoir’s surface, respectively). Finally, low risk areas covered 27940 and 25996 ha, for cyanobacteria abundance and chlorophyll-a concentration, respectively (58-62% of the reservoir’s surface, respectively).

**Figure 5.**
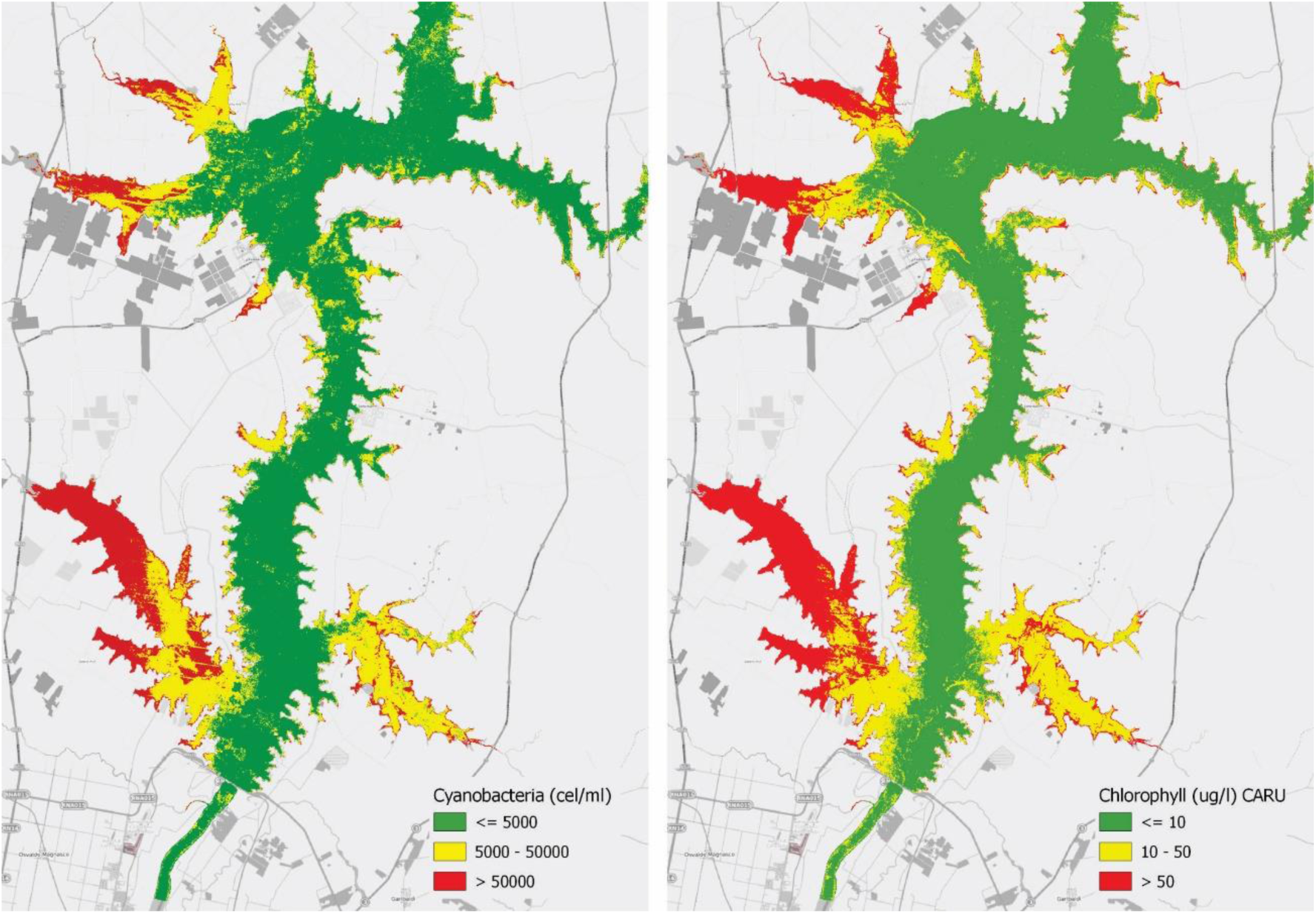
Mean distribution of cyanobacteria abundance (left) and chlorophyll-a concentration (right) for the warm months (December to March) of the years 2011-2017 following the risk categories established by the Comisión Administradora del Río Uruguay (CARU) for recreational use of waters.

When comparing CARU with WHO guidelines for cyanobacteria, both guidelines showed similar spatial patterns (Supplementary Material 4). As expected, CARU, which has more conservative thresholds, rendered greater areas within the high and moderate risk categories, yet fewer areas within the moderate risk category, compared to WHO (high risk: 13% CARU, 7% WHO, moderate risk: 25% CARU, 14 % WHO, low risk: 62% CARU, 78% WHO).

The PCA explained 95% of total variability (first axis explained 62.5% and the second axis 32.2%). Chlorophyll-a measured with satellites, chlorophyll-a measured in the field, and cyanobacteria abundance estimated with satellites were closely located in the PCA (Fig. 6). Also, cyanobacteria measured in the field and the cyano-traffic-light reports were positively closely located in the PCA

**Figure 6.**
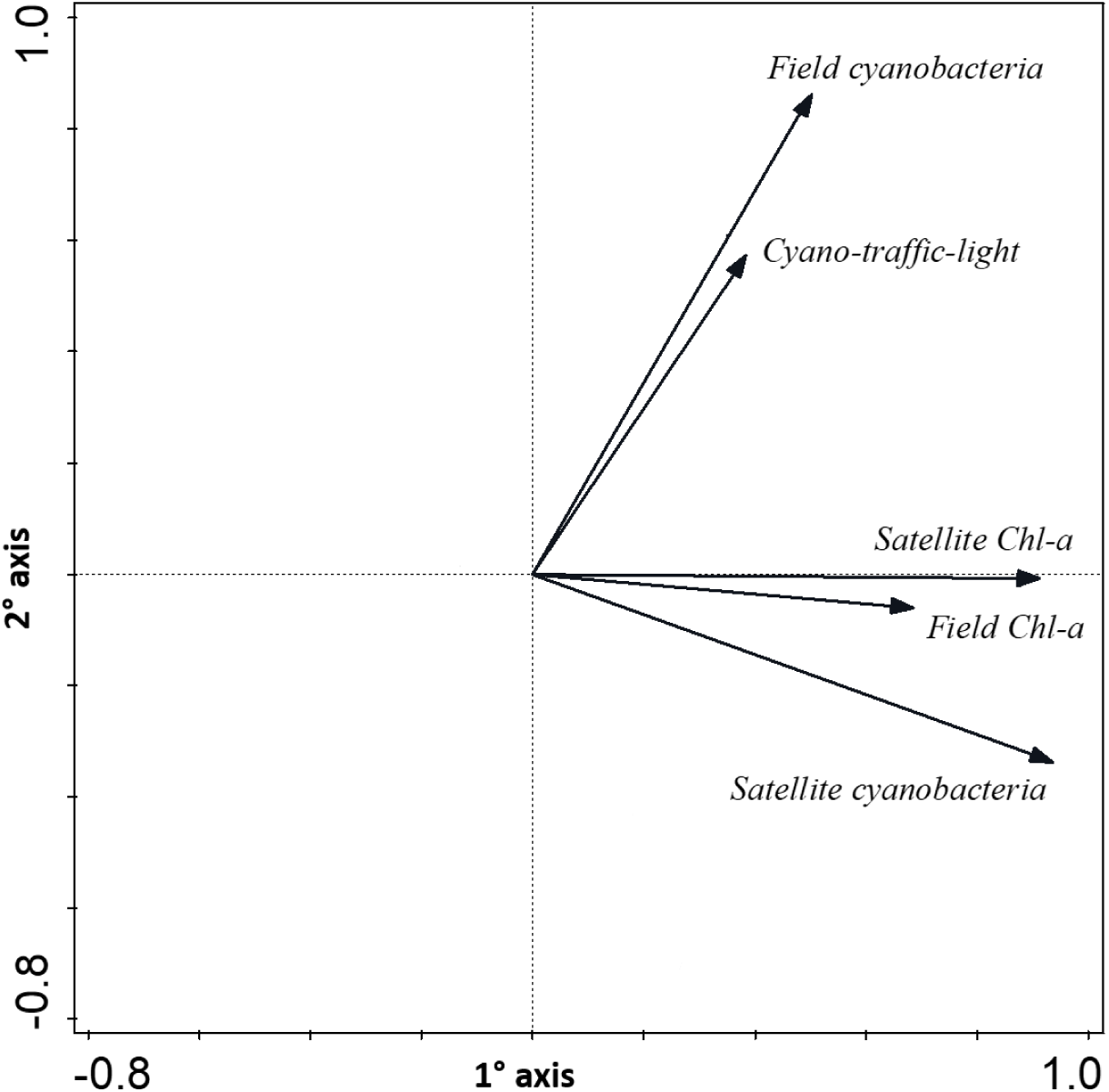
Principal component analysis using the variables obtained in the field and estimated with satellite images.

(Fig. 6). Nevertheless, cyanobacteria measured in the field (and the cyano-traffic-light reports) and cyanobacteria measured with satellites (and field and satellite chlorophyll-a) lacked correlation (Fig. 6).

### Assessment of cases of possible human intoxication due to recreational exposure to hazardous bacteria

From 8^th^ January to 8^th^ February 2017 a total of 39 surveys were completed, as these patients evidenced symptoms that could be related to hazardous bacteria intoxication. Out of 39 surveys, 20 had recreationally used waters form the Uruguay River. Out of these 20 surveys, only 8 exposed to waters in the Salto Grande Reservoir (study area); the remaining 12 suspected cases recreationally used waters located outside of the study area (down waters from the reservoir) and hence were not considered for further analyses. For the 8 cases recorded in the Salto Grande reservoir, the most frequent symptoms were diarrhea and vomits, followed by otitis and few cases of rash (Table 4). Most patients used recreational sites for bathing, from about 30 minutes to up to two hours; the time span between the recreational exposure and the visit to the hospital was mostly of one to two days (median=2 days) (Table 4). The suspected cases occurred in all 4 recreational areas located in the southern part of the Salto Grande reservoir (Concordia department) (Table 4, Fig. 7), and suspected cases were reported every 2 to 6 days (Table 4). Out of the 4 sites with suspected cases, Sol and Los Medicos had high-risk of coliform bacteria but low risk of cyanobacteria, site Las Perdices had high risk of coliform bacteria but moderate risk of cyanobacteria, while site Las Palmeras had high-risk of both hazardous bacteria (Table 4, Fig. 7).

**Figure 7.**
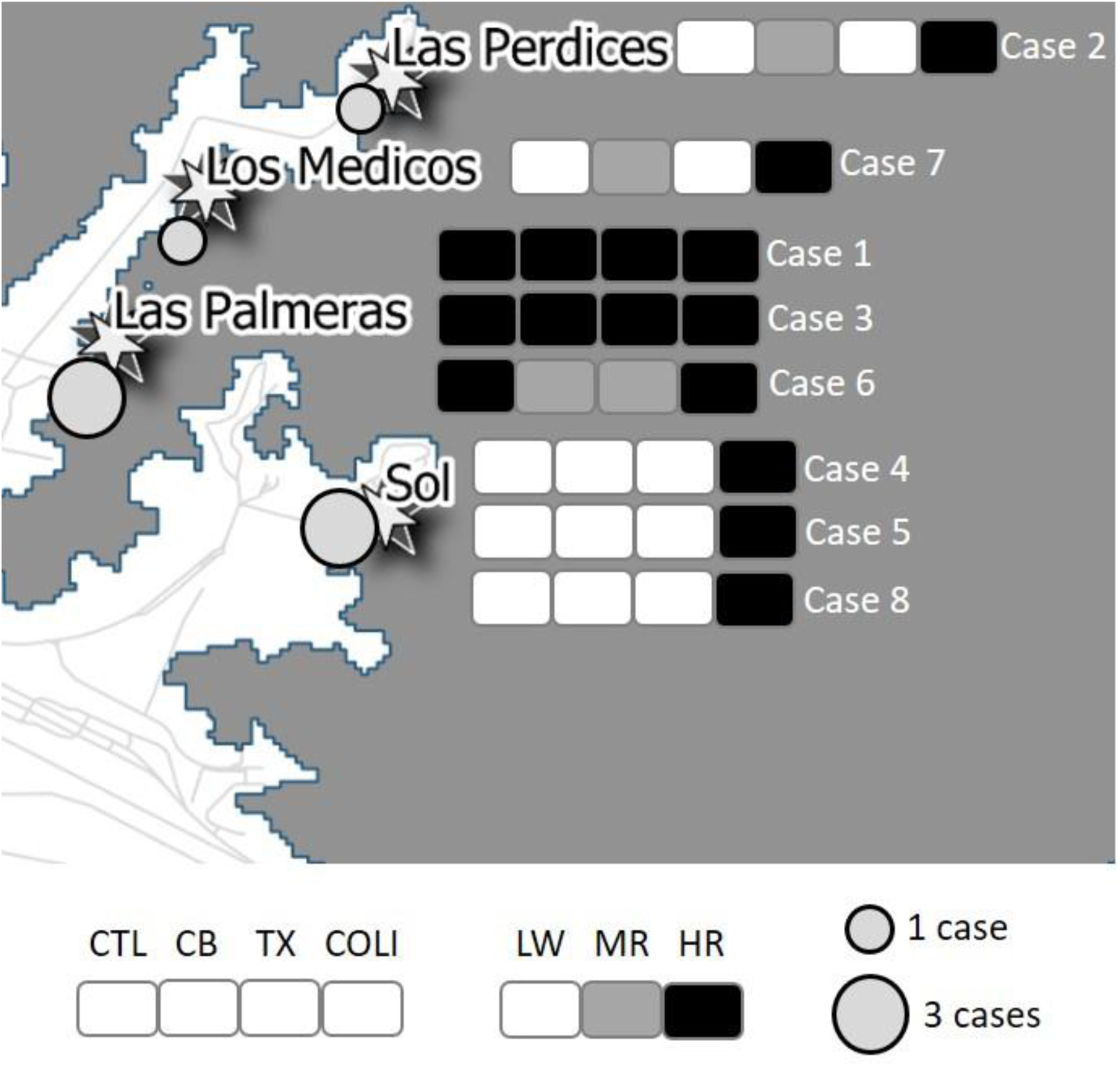
Environmental risk posed by hazardous bacteria in the recreational areas located in the southern part of the Salto Grande reservoir, for the dates where the 8 possible cases were identified. Case numbers are detailed in Table 4. CTL=Cyano-traffic-light, CB=cyanobacteria abundance, TX=microcystin concentration, COLI=coliform bacteria concentration.

**Table 4.**
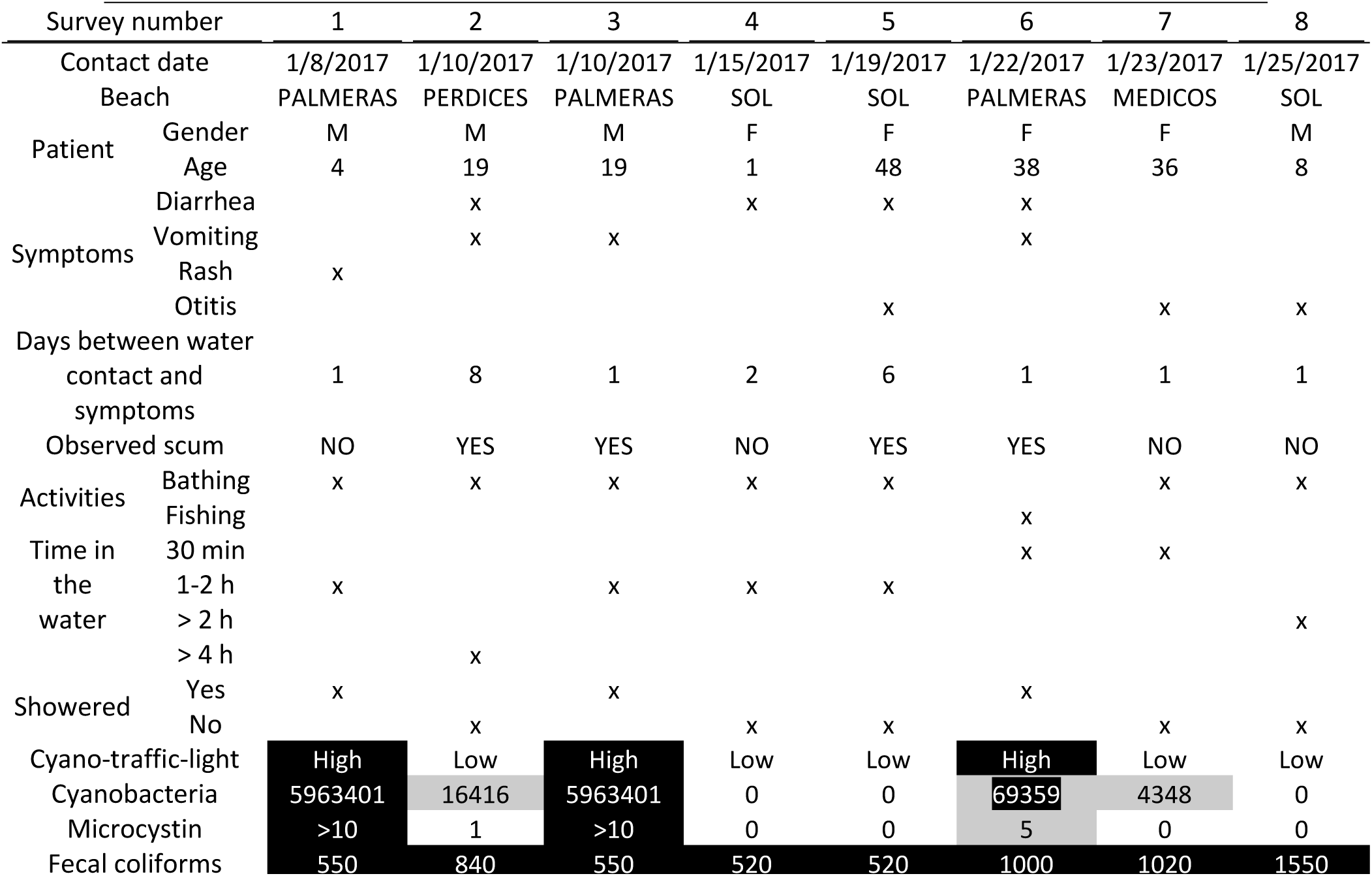
Results of the surveys for the suspected cases of intoxication with hazardous bacteria and that had recreational contact with the Salto Grande Reservoiŕs water. For cyanobacteria abundance (cell/ml) and microcystin concentration (µg/l) the risk categories (low, medium, high) are informed. For fecal coliforms (CFU/100 ml), when the threshold of 500 CFU100 mL^-1^ was exceeded, the risk was considered high. White: low risk, Gray: moderate risk, Black: high risk.

## Discussion

Our results show that in summer, when water use for recreation is highest in the Salto Grande reservoir, the environmental risk posed by coliform bacteria and cyanobacteria was high, and that cases of suspected illness due to recreational exposure were identified. Bellow, we discuss the main findings, strengths, and limitations within and among the components (field monitoring, remote sensing monitoring and health survey) analyzed.

### Assessment of the spatial and temporal distribution of cyanobacteria and coliform bacteria: water monitoring at recreational areas

The field study allowed detailed information of cyanobacteria (abundance and toxicity) and *E. coli* concentrations, yet with a limited spatial (about 50% of official recreational sites) and rather low temporal (four years) coverage. Cyanobacteria, abundances recurrently surpassed the moderate and high-risk thresholds established by CARU across sites and year analyzed. A positive and significant link between cyanobacteria abundance and microcystin concentration was found, as observed elsewhere (Pearl and Otten 2013; Ibelings et al. 2014). Nevertheless, microcystin concentrations were mostly within the low to moderate risk categories established by CARU (<10 µg L^-1^), though on few occasions (10%) they were within the high-risk category (>10 µg L^-1^). Other studies in the same reservoir also found microcystin concentrations in the high-risk category (Martinez de la Escalera et al. 2017, maximum value recorded:15 µg L^-1^; Bordet et al. 2017, maximum value recorded: 621 µg L^-1^), underscoring the potential harm that toxins can cause to health in the studied area.

Mean coliform bacteria was within the high-risk category in both recreational areas located in the north part of the reservoir (Federación department). It is probable that the sewage system of the city of Federación is probably inefficient, mostly during summertime when its population increases 400 % due to tourism. Indeed, a report produced by CARU (2016) recognizes that only few localities along the coastline of the lower stretch of the Uruguay river have the sufficient treatment facilities to respond to their number of inhabitants. Recreational areas located in Concordia department, however, were within the low-risk category for *E. coli*, probably because these sites are about 15 km north from the closest city (Concordia). Nevertheless, in 2017 *E. coli* values in these sites were in the high-risk category (see below). We found absence of a relationship between cyanobacteria (cell abundance and microcystin) and coliform bacteria abundances, probably because these hazardous bacteria respond to different environmental triggers. For example, cyanobacteria (photosynthetic organisms) development is promoted by high temperature, long photoperiod, and low wind (Paerl and Huisman, 2009), whereas *E. coli* (heterotrophic bacteria) is mostly associated to fecal sources.

### Assessment of the spatial and temporal distribution of the risk posed by cyanobacteria at landscape scale: remote sensor monitoring

The use of satellite images allowed describing the extent and intensity of the cyanobacteria blooms at landscape scale (760km^2^), integrating interannual variations throughout the growing season of a 7-year period (2011-2017). Hence expanding the temporal and spatial scale characterized in the field study. The resulting risk map provides baseline information for the spatial distribution of the risk posed by cyanobacteria both for Argentina and Uruguay, which can help water managers to effectively distribute resources to monitor and manage its waters. The emerging picture is that both cyanobacteria abundance and chlorophyll-a have low risk in the river like area, moderate to high risk in the Argentinean arms, and moderate to low risk in the Uruguayan arms. As stated in the introduction winds concentrate blooms towards the Argentinean coast (O’Farrell et al. 2012; Bordet et al. 2017). Interestingly, our results at landscape scale are consistent with patterns obtained in field studies in this reservoir (CARU 201, Bordet et al. 2017). Noteworthy, areas identified as having the highest risk remain currently unmonitored by CARU and CTM water agencies.

The use of satellite images allowed bridging the gaps of information and the limited spatial coverage provided by field samplings. Nevertheless, compared to field studies, satellites are unable to monitor toxicity nor fecal bacteria, as these lack optical properties. Cyanotoxins could be estimated using indirect proxies such as chlorophyll-a estimated with satellites, as done by Shi et al. (2015). Also, based upon the positive link between cells and toxicity observed in our and other studies (Pearl and Otten 2013; Ibelings et al. 2014), areas with high risk due to cyanobacteria abundance could be explicitly considered as high toxicity too. Future field monitoring in the arm areas (which are currently mostly unmonitored) is needed for assess how representative the information on cyanobacteria abundance is of toxicity. Finally, this baseline risk map can be coupled with several vulnerability variables, such as access to coastline, inhabitants in cities and villages, availability of sewage facilities, use of water, among other variables to use as input for assessing possible health risks.

We found comparable results for field and satellite results in terms of chlorophyll-a but not for cyanobacteria abundance. The latter differences probably aroused from comparing different metrics with different scales (point sampling in the field versus sampling areas with satellites) and sampling methods (integrated samples in the field versus superficial monitoring with satellites) and monitoring frequencies. Indeed, comparing results across a diverse set of metrics can be challenging (Ho and Michalak, 2015). Nevertheless, as mentioned before, we found a similar distribution of cyanobacteria risk both in field and in satellite approaches.

Welker et al. (2021) suggested that remote sensing data could be integrated into early warning systems to protect human health and ecosystems. Indeed, in the Salto Grande reservoir, cyanobacteria abundance monitored with satellites could be used for determining the risk categories of the cyano-traffic-light, at least for sites that currently remain unmonitored (Table 5). The near real-time surveillance of cyanobacteria blooms by satellites provides a rather low-cost opportunity to intensify monitoring in time and space (Welker et al. 2021), particularly in this reservoir where large areas identified as high-risk using satellites remain unmonitored by CTM and CARU water agencies. We, however, suggest excluding chlorophyll-a monitored by satellites from the cyano-traffic-light, as this variable is common to all phytoplankton and could lead to overestimations or underestimations of the risk posed by cyanobacteria.

**Table 5.** Suggestions of improvement to the criteria used by the Comisión Administradora del Río Uruguay (CARU) for establishing the risk categories of the cyano-traffic-light for recreative use of water

| Proxy | Low Risk | Moderate Risk | High Risk |
| --- | --- | --- | --- |
| Cyanobacteria (cell mL<sup>-1</sup>)* | <20,000 | 20,000-100,000 | >100,000 |
| Microcystin (µg L<sup>-1</sup>) | < 24 | ≥24 | > 24 |
| *E.coli* (CFU 100 mL<sup>-1</sup>) | <500 | 500-900 | >900 |
| Cyano-traffic-light color | Green | Yellow | Red |
| Bathing recommendation | Allowed | Avoid using | Prohibition of use |

Future improvements for increasing the comparability between remote sensing and field metrics, include: i) planning field samplings in dates that match the passing of satellites and iii) include Sentinel imagery (launched after the ending of our study) which have better temporal, spatial, and spectral resolution than Landsat imagery, and which can differentiate cyanobacteria from other phytoplankton. We already have algorithms for measuring cyanobacteria abundance using Sentinel imaginery in Salto Grande Reservoir (Drozd et al. 2019). Average growth rate for cyanobacteria is of 0.4 days^-1^ (Schwaderer et al., 2011), which suggests that doubling time takes about two and a half days to occur. Hence the joint use of different satellites, such as Landsat and Sentinel, would allow a monitoring frequency of days that would allow tracking well cyanobacteria population growth rates.

### Assessment of possible cases of human intoxication due to recreational exposure to hazardous bacteria

The number of surveys obtained for assessing the link between recreational exposure to hazardous bacteria and illness at Salto Grande Reservoir was very low to draw any conclusion. Nevertheless, we would like to highlight several findings that can contribute to the scarce literature of health effects associated to recreational exposure to cyanobacteria (Stewart et al. 2006b; Wood 2016) and pathogenic organisms in ambient water (Russo et al. 2020). We detected cases with symptoms in four out of the six recreational sites analyzed. These four sites are located 15km away from the hospital where the survey was performed, in the southern part of the reservoir. The absence of cases in the two recreational sites located in the northern part of the reservoir (Federación City) seem more related to the large distance between cities –about one hour commuting– and to the fact that Federación has its own hospital (patients with symptoms would visit its own hospital), than to absence of risk in these sites (which in the field study evidenced high risk of fecal bacteria) or absence of symptoms in end users. Patients with symptoms and who recreationally exposed to water belonged to Concordia city (where the hospital was located) were of a wide range of age and gender, and evidenced mostly gastrointestinal (vomiting, diarrhea), and ear (otitis), and to a lesser extent dermal (rash) symptom. A fairly short time lag occurred between exposure and symptom appearance: with symptoms appearing mostly within one to two days after exposure. Most patients showed a high degree of exposure to recreational waters as they reported swimming for up to two hours. Noteworthy, several of the health cases that occurred outside from the studied area (down waters from the Salto Grande reservoir) seemed related to high coliform bacteria concentrations (cyanobacteria abundances were low) (data not shown).

Despite the overall low number of cases, these were identified with high temporal frequency (every 2 to 5 days within a month). Wood (2016) also found that the number of cases of intoxications due to recreational exposure to cyanobacteria affected few numbers of people yet occurred with high recurrence. The low number of cases obtained in our study can relate to several reasons: 1) symptoms are mild, short lasted and self-limiting (Steward et al. 2006b), and this may result in under-reporting of symptoms by affected patients, 2) patients may have visited other medical facilities in Concordia city or other cities 3) low compromise of physicians and health assistants (see below) in recording the surveys, probably due to the high demands of their work.

In the different recreational areas with suspected health cases, the risk posed by hazardous bacteria at the time of exposure was mostly related to coliforms (beaches Sol, Las Palmeras, Los Medicos, Las Perdices) and to a lesser extent to cyanobacteria (mostly in beach Las Palmeras, that was the only site with high toxin risk). Nevertheless, when allocating symptoms to bacteria hazards in water it is important to realize that mere co-occurrence is insufficient for establishing causal connection (Chorus and Welker, 2021). Noteworthy, at the onset of our study, we expected to find higher risk due to cyanobacteria than to coliforms, probably because cyanobacteria blooms are conspicuous (due to their pigments they can turn the water green) in the reservoir. In this line of thought, end users need to be informed about the risk they are exposed when using water recreationally (they can actively choose to avoid green waters due to high cyanobacteria abundance, but bath unaware of risk if coliforms which are colorless), to take informed decisions and avoid illness.

The risk posed by fecal coliforms seems to have increased over time: in the field approach run during 2011-2015, risk posed by mean coliform bacteria was low, whereas in summer 2017 (when the survey was run) the same sites presented high risk of fecal coliforms. This result once again reinforces the urgent need to incorporate the measurement of coliform bacteria into the risk communication system for recreational exposure to water in the Uruguay river. It also underscores the need of undertaking actions towards decreasing the causes that originate high concentrations of fecal coliforms, such as improving sewage systems where necessary.

Finally, the fact that we found suspected cases, and that isolated cases have been identified in the past (Giannuzzi et al. 2011), denotes situations where aspects of the biophysical environment affect human health. It also underscores the need to continue monitoring suspected human health cases linked to recreational exposure in this ecosystem.

Despite the evidence of the direct and indirect effects of hazardous bacteria on public health, environment professionals, water managers and public health professionals rarely interact. Our attempt to work interdisciplinary faced its highest challenges when interacting with health professionals. Despite the head of the physicians and health assistants showed a high interest in undertaking this study, and collaborated promptly at the onset of the project, then they were unreachable during the follow up process (despite many attempts from our part). Moreover, the number of surveys collected was too low compared to the number of patients that visit the emergency room, which shows a rather low compromise of action in undertaking this endeavor. Indeed, our work showed a high degree of integration among academia (ecologists, remote sensing experts) and water management agencies, yet a poor integration with health experts (despite many attempts to interact). Likewise, El Saadi et al. (1995) alluded to the difficulty of gaining the cooperation of medical practitioners when assessing health cases related to recreational exposure to cyanobacteria. The latter barrier needs to be surpassed to identify how exposure to water born microorganisms relate to health effects, with the aim of protecting humans from illness.

### Assessment of the effectiveness of the risk communication system and improvement suggestions

The risk communication system developed by CARU for cyanobacteria (cyano-traffic-light qualitative reports) evidenced an overall similar pattern to cyanobacteria mean abundance, mostly in the year of highest cyanobacteria abundance (2012). Noteworthy, the cyano-traffic-light is built upon the surpassing of at least two of the four variables used by CARU to establish the different risk categories (cell abundance, toxicity, scum, and chlorophyll-a). Hence even if cyanobacteria abundances may indicate high risk, if the other 3 variables indicate a low risk, the observed cyano-traffic-light would indicate a low risk. The predictive model was significant only when using cyanobacteria abundance as predictive variable, probably because we had plenty of data for this variable, but few data for microcystin concentration and even less data for chlorophyll-a. When comparing the observed with the predicted cyano-traffic-light, the explanatory power was low, with the predicted cyano-traffic-light overestimating the observed cyano-traffic-light. As mentioned above, this probably occurred because at least two proxies are used to determine the outcome of the cyano-traffic-light, while the statistical model only used cyanobacteria abundance.

After almost a decade of using the cyano-traffic-light, and based upon our observations, we suggest several improvements to this alert system (Table 5): 1) using toxicity and cyanobacteria abundance but excluding chlorophyll-a and presence of scums. Microcystin, is the most reported toxin worldwide (WHO 2020) in health relevant concentrations (Svirčev et al. 2019) and is the only cyanotoxin monitored in the Salto Grande reservoir. This approach is adequate as the dominant (*Microcystis*) and subdominant (*Dolichospermum*) cyanobacteria that bloom in the Salto Grande reservoir (Bordet et al. 2017) are microcystin producers. Nevertheless, because species within *Dolichospermum* can also produce anatoxins (ANA) and saxitoxins (SAX) (Li et al. 2016 we), we encourage the incorporation of the assessment of these toxins whenever *Dolichospermum* species are detected. The later would decrease the current uncertainty about these toxins in the reservoir and would allow assessing the need of including them (or not) into the cyano-traffic-light variables. Regarding cyanobacteria abundance, keeping this proxy in the building of the cyano-traffic-light, would allow protecting the population against its toxins and against unspecific health effects of cyanobacteria blooms not attributable to cyanotoxins (Chorus and Welker, 2021). Indeed, cyanobacteria blooms are often caused by heavy nutrient loads, which can be associated with loads of pesticides and/or other contaminants from agriculture and/or poorly treated wastewater and/or organisms that may be hazardous (Chorus and Welker, 2021). Hence, it is therefore prudent to avoid exposure to high concentrations of cyanobacteria even if concentrations of toxins are low (Chorus and Welker, 2021). In the revision of the guidelines used for cyanobacteria risk assessment, it has been proposed that biovolume should be used instead of cell numbers, to avoid overestimation of cell abundances and toxicity when small celled sized cyanobacteria species dominate (Chorus and Welker, op cit.). Nevertheless, we suggest keeping the use of cell numbers in the cyano-traffic-light, because: i) cyanobacteria species dominating blooms in the reservoir are rather big sized and have similar cell diameters (range: 5-10µm) and ii) the use of cell numbers would allow comparing field and satellite approaches (see next section). We suggest removing chlorophyll-a from the cyano-traffic-light as it can either overestimate (at periods without cyanobacteria dominance or in mixed blooms of cyanobacteria and eukaryotic phytoplankton) or sub estimate (when dense cyanobacteria blooms have low chlorophyll-a concentration) the risk posed by cyanobacteria abundances. We also suggest removing the proxy of scum formation from is markedly affected by wind action and its absence may erroneously indicate absence of high densities of cyanobacteria. 2) Moreover, for cyanobacteria abundance and toxin concentration, we suggest increasing the concentrations that determine the thresholds of each risk category, following the values proposed of Chorus and Bartram (1999) for cyanobacteria abundance and Chorus and Welker (2021) for microcystins (Table 5). 3) We suggest the inclusion of *E. coli* in the cyano-traffic-light (Table 5). The proposed revised version of the cyano-traffic-light can be named bacteria-traffic-light, as it would include the risk of two of the several bacteria that can trigger illness in recreational waters. 4) Finally, we suggest that for allocating of a risk category in the cyano-traffic-light at least one variable-cyanobacteria, toxicity or *E.coli*-should be surpassed (currently at least two variables are computed).

## Conclusion

Our study evidences the advantages of integrating disciplines (ecology, water management, health) and metrics (quantitative and qualitative field data, remote sensing data) in characterizing the risk posed by water born bacteria in recreational waters, capitalizing on the strengths, and minimizing the weakness of each individual component. Finally, as in many other ecosystems around the world, the Salto Grande reservoir is very well studied from the environment standpoint, yet there is almost absence of studies assessing its link to human health. Bridging this gap requires the integration among environmental management and public health disciplines.

## Acknowledgements

We are grateful to the monitoring agencies Comisión Administradora del Río Uruguay (CARU) and Comisión Técnica Mixta (CTM), and their working teams, in particular Natalia Rougier, for facilitating field data, and Pilar Ojeda, Hector Procura, Jorge Blasig, Heleno Zabaleta, Ricardo Juárez y Fernando Gauna for data field collection used on sattelite models building and validation. We are thankful to the Hospital Masvernat and the health group headed by Dr. Guillermo Saucedo for participating in the survey study. We thank the Ethics committee from Garrahan Hospital for performing the ethics evaluation for the survey. We are thankful to the Área de Investigación Científica y Tecnológica of the Faculty Exact and Natural Science, University of Buenos Aires (FCEyN-UBA), for their help in establishing the information transfer agreement between the Masvernat Hospital and FCEyN-UBA. We are thankful to Tatiana Petcheneshsky for all her support during the preparation of the health study. DG acknowledges CARU scholarship, and Oñativia scholarship from the National Argentinean Health Ministry.

---

**Supplementary Material 1.**
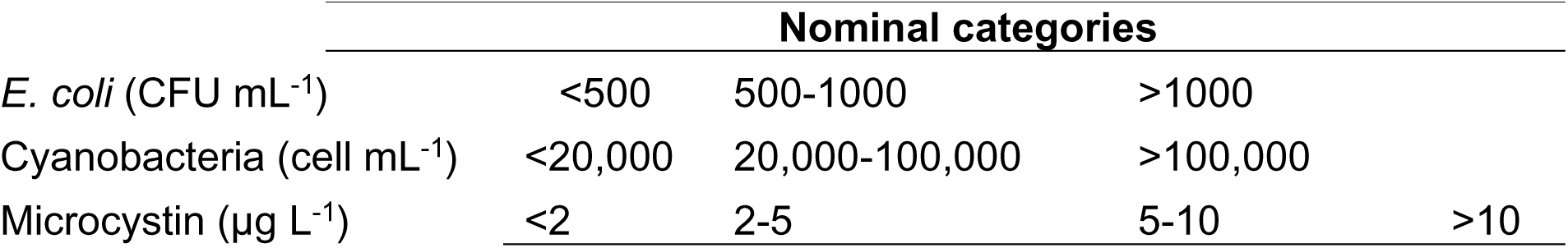
Categories used for the contingency analysis.

**Supplementary Material 2.**
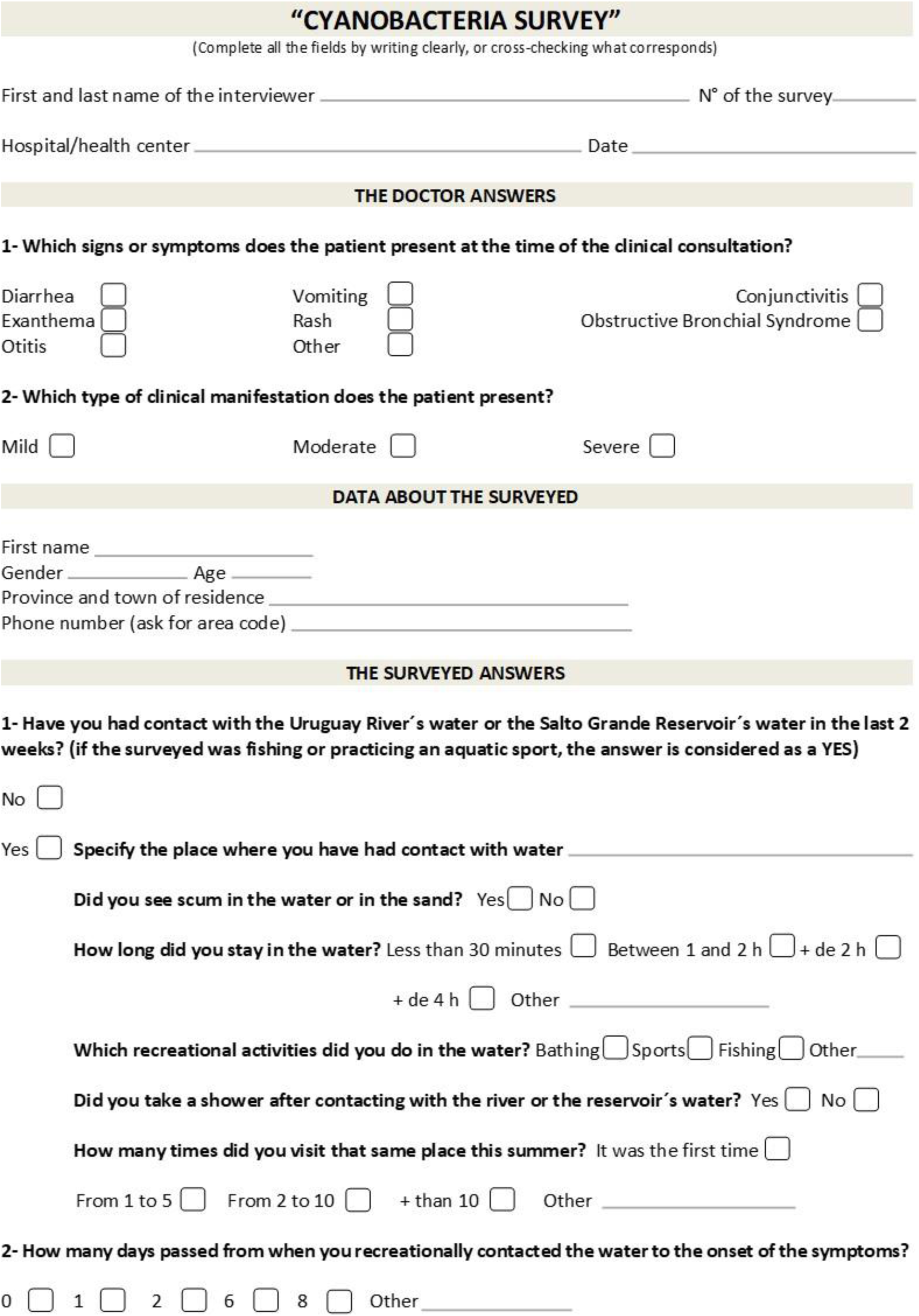
Survey performed at the Masvernat Hospital in summer 2017

**Supplementary Material 3.**
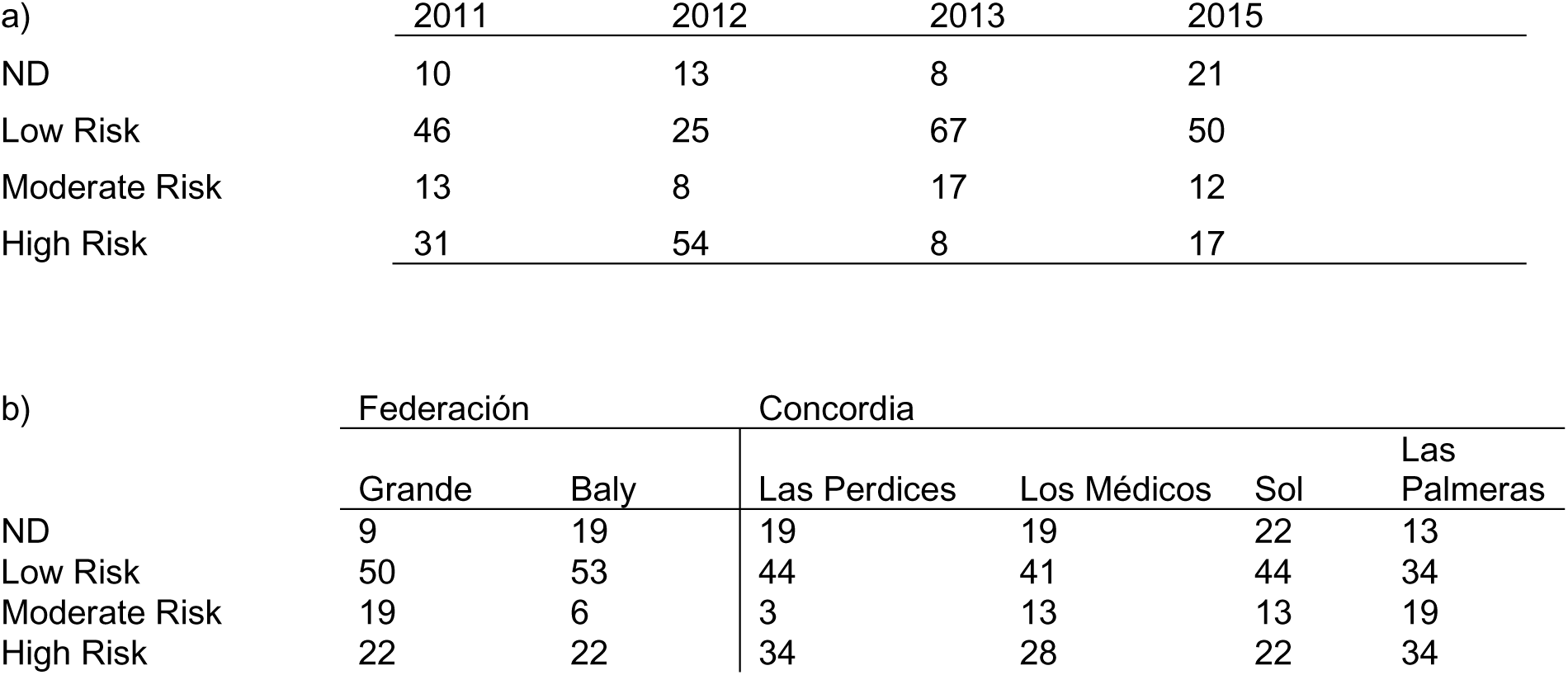
Mean percentage of the cyano-traffic-light categories registered by a) warm season of each year and b) each recreational area. ND= no data.

**Supplementary material 4.**
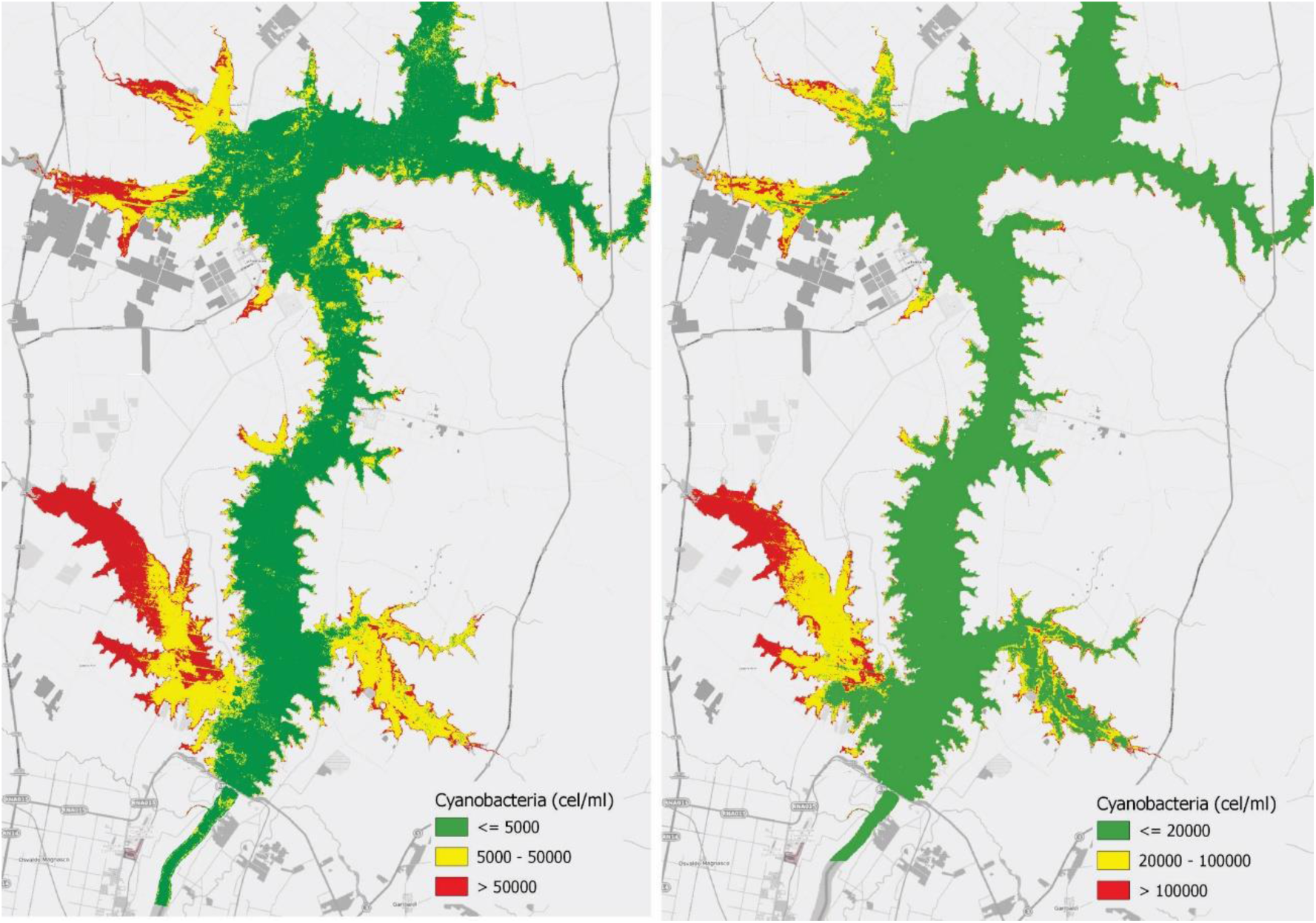
Distribution of risk posed by mean cyanobacteria abundance in the Salto Grande Reservoir (warm periods of 2011-2017), sorted into the CARU (Comisión Administrativa del Río Uruguay) (left) and WHO (World Health Organization) (right) risk categories.

